# Genetic Diversity and Distributional Pattern of Ammonia Oxidizing Archaea Lineages in the Global Oceans

**DOI:** 10.1101/249441

**Authors:** Shunyan Cheung, Wingkwan Mak, Xiaomin Xia, Yanhong Lu, Hongbin Liu

## Abstract

In the study, we used miTAG approach to analyse the distributional pattern of the ammonium oxidizing archaea (AOA) lineages in the global oceans using the metagenomics datasets of the *Tara* Oceans global expedition (2009-2013). Using ammonium monooxygenase alpha subunit gene as biomarker, the AOA communities were obviously segregated with water depth, except the upwelling regions. Besides, the AOA communities in the euphotic zones are more heterogeneous than in the mesopelagic zones (MPZs). Overall, water column A clade (WCA) distributes more evenly and widely in the euphotic zone and MPZs, while water column B clade (WCB) and SCM-like clade mainly distribute in MPZ and high latitude waters, respectively. At fine-scale genetic diversity, SCM1-like and 2 WCA subclades showed distinctive niche separation of distributional pattern. The AOA subclades were further divided into ecological significant taxonomic units (ESTUs), which were delineated from the distribution pattern of their corresponding subclades. For examples, ESTUs of WCA have different correlation with depth, nitrate to silicate ratio and salinity; SCM1-like-A was negatively correlated with irradiation; the other SCM-like ESTUs preferred low temperature and high nutrient conditions, etc. Our study provides new insight to the genetic diversity of AOA in global scale and its connections with environmental factors.

## Introduction

Nitrification is an important and well-concerned pathway in the marine nitrogen cycle, which can supply approximately 25-36 % of bioavailable nitrogen to phytoplankton (Wankel et al., 2007; Yool et al., 2007). Besides that, it was also estimated as dominant source of nitrous oxide (Freing et al., 2012), a strong greenhouse gas influencing the Earth climate (Solomon, 2007). As the first and rate-limiting step of nitrification, ammonia oxidation was formerly only discovered within bacterial domain (AOB) (Ward, 2002). However, unique archaeal-associated ammonia monooxygenase gene recovered from environmental metagenomics data over a decade ago, suggested the metabolic potential of archaeal ammonia oxidation (Venter et al., 2004; Treusch et al., 2005). Using ammonia monooxygenase alpha subunit (*amoA*) as a biomarker, archaeal *amoA* genes were detected throughout water column and marine sediments (Francis et al., 2005). With the first isolation of marine ammonia oxidizing archaea (AOA) (*Nitrosopumilis maritimus* SCM-1), AOA are known to be chemo-lithoautotrophor, aerobically oxidizing ammonia to nitrite while fixing inorganic carbon (Könneke et al., 2005). Subsequent field studies revealed AOA outnumber AOB in the oceanic waters (Wuchter et al., 2006; Mincer et al., 2007; Beman et al., 2008; Beman et al., 2010; Santoro et al., 2010), and they play a dominant role in determining the activity and distribution of nitrification process (Wuchter et al., 2006; Beman et al., 2008; Smith et al., 2014). In contrast to AOB, AOA (*N. maritimus* SCM-1) exhibited exceptionally high ammonia affinity and low thresholds, explaining why AOA are dominant in the oceanic waters (Martens-Habbena et al., 2009).

In the first phylogenetic analysis of AOA *amoA* gene, two groups of AOA were commonly detected in the water column, and they were named water column A (WCA) and water column B (WCB), respectively (Francis et al., 2005). WCA and WCB are considered as the major ecotypes of AOA in the oceanic waters, predominating in upper (< 200m) and deep (> 200m) layers, respectively (Francis et al., 2005; Beman et al., 2008; Santoro et al., 2017). So far, there is only one culture of WCA (*Nitrosopelagicus brevis* CN25) reported (Santoro and Casciotti, 2011; Santoro et al., 2015) and the WCB is still uncultivated, therefore, the physiology of these two ecotypes are still unclear (Santoro et al., 2017). In spite of that, several factors have been hypothesized as potentially causing the vertical succession of WCA and WCB, including light inhibition, differential affinity and utilization of nitrogen sources, and different requirement of micro-nutrients (Amin et al., 2013; Sintes et al., 2013; Luo et al., 2014; Qin et al., 2014; Sintes et al., 2016; Smith et al., 2016; Santoro et al., 2017).

With development of DNA sequencing technique, our knowledge of the genetic diversity of AOA is improving (Sintes et al., 2016; Jing et al., 2017). Using next generation sequencing to samples from western subarctic Pacific Ocean, diverse subclades of ecotype WCA and WCB were defined; and some subclades of same ecotypes showed differential distributional patterns (Jing et al., 2017). In addition to that, recently global scale study has found that the ecological significant taxonomic units (ESTUs) of *Synechococcus* were delineated from their corresponding ecotypes, in which OTUs of same ecotypes were grouped according to their distributional patterns (Farrant et al., 2016). Therefore, although most of the previous studies were targeting the whole clade of WCA and WCB (Francis et al., 2005; Beman et al., 2008; Santoro et al., 2010; Smith et al., 2014; Smith et al., 2016; Santoro et al., 2017), the subclades and ESTUs of WCA and WCB may show distinctive distributional patterns in the oceans. Moreover, analysing the subclades and ESTUs of AOA in diverse marine ecosystems may provide better and more insights on the specific connection between genetic diversity of AOA and environmental conditions.

In this study, we conducted detailed phylogenetic analysis of AOA in the global oceans using metagenomics datasets of *Tara* Oceans expedition (2009-2013) (Karsenti et al., 2011; Armbrust and Palumbi, 2015). We have built a database of *amoA* gene sequences and recovered the AOA communities from the metagenomics datasets using miTAG approach (Logares et al., 2014). miTAG is an approach mapping short reads of metagenomics dataset to long sequences of known identity, which has been proven as a promising and PCR amplification biases free method for recovering community of bacteria and picocyanobacteria from the *Tara* Ocean datasets (Logares et al., 2014; Farrant et al., 2016). Furthermore, we have further divided the subclades of AOA into ecological significant taxonomic units (ESTUs), in which OTUs of same subclades were grouped according to their distributional patterns (Farrant et al., 2016). Through all these analyses, we described the distributional patterns of AOA and characterized the AOA ESTUs with environmental conditions.

## Methods

### Constructing database of *amoA* gene

In order to retrieve and identify the *amoA* affiliated sequences (AARs) in the *Tara* Oceans metagenomics dataset (Karsenti et al., 2011; Armbrust and Palumbi, 2015), the reference sequences and environmental sequences of *amoA* gene were downloaded from National Centre for Biotechnology Information (NCBI) nucleotide database (Acland et al., 2014). These sequences were aligned with MUSCLE (Edgar, 2004) and the majority of sequences have overlapping at the region targeted by the Arch-amoAF/Arch-amoAR primer set (Francis et al., 2005). The sequences that have overlapping in this region were chosen and clipped into an alignment with the same length, and then were used to calculated OTUs at 97 % DNA similarity cut off using Mothur (Schloss et al., 2009). Besides that, amino acid sequences of these selected sequences were also downloaded from the NCBI (Acland et al., 2014).

### Recruiting AARs from the *Tara* Oceans dataset

The clean dataset of *Tara* Ocean metagenomics dataset (processed nucleotide reads, 100-200 bases per reads) were downloaded from the European Bioinformatics Institute (EMBL-EMI) (https://www.ebi.ac.uk/metagenomics/projects/ERP001736) and aligned against the amino acid sequences of our *amoA* gene database using DIAMOND (e-value = 1e^-5^) (Buchfink et al., 2015). DIAMOND is a BLASTX like algorithm 2000 times faster than BLASTX on short reads (Buchfink et al., 2015), which reduces the demand of computational resources when recruiting the *amoA* from large metagenomics dataset of the *Tara* Oceans. The sequences recruited with DIAMOND were then blasted against the DNA sequences of our *amoA* database using BLASTN (McGinnis and Madden, 2004). The reads which have 99-100 % similarity and coverage with the DNA sequences in our *amoA* database were chosen as AARs. There was less than 1 % of reads showed lower than 99 % similarity with the sequences in our dataset, which were neglected in downstream analysis. AARs were recruited from 35 bacterial size fraction metagenomics datasets that contained significant number of AARs. Except 3 datasets, which contains 35, 43 and 51 AARs, the rest datasets contained more than 100 AARs each (111-10,604 reads) (S 1 Table). These datasets contain 22 and 13 samples from euphotic zone and MPZ, covering Pacific Ocean, Atlantic Ocean, Indian Ocean, Southern Ocean, Red Sea, Mediterranean Sea and Gulf of Mexico (Fig. 1).

**Fig. 1.**
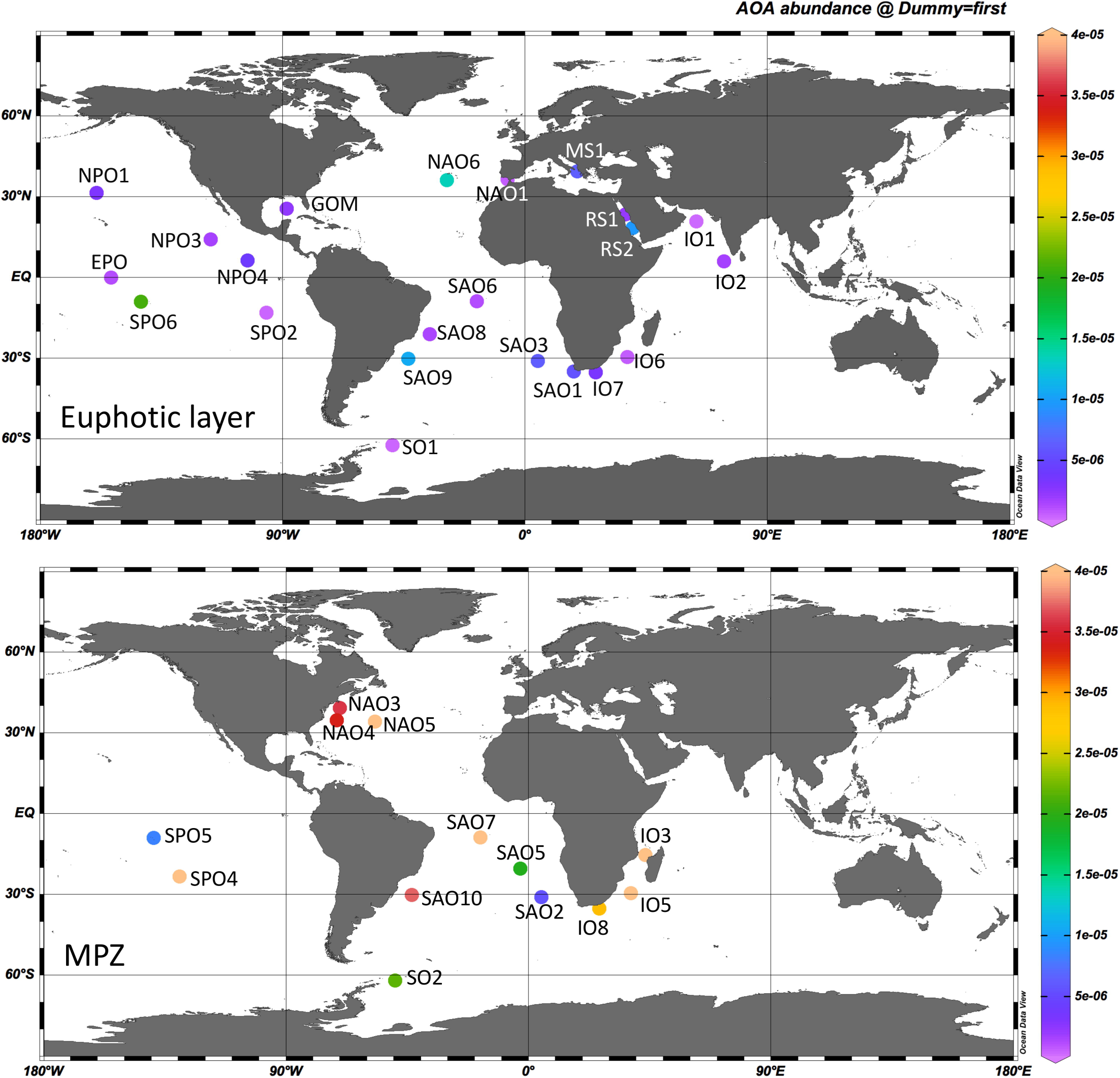
Sampling locations of metagenomics datasets selected in this study. The relative abundances of AOA (AARs divided by total clean reads) in the metagenomics datasets were shown with color gradient.

### Phylogenetic analysis

The number of AARs which affiliated to the DNA sequences of the same OTUs were merged into the sequence number of each OTUs; and the relative abundance of each OTU in each sample were calculated by dividing sequence number of the OTU in particular sample by total clear reads of that sample. Because the number of AARs in different samples are highly varied, ordering of the OTUs were normalized by their relative abundance, instead of sequence numbers. Top 202 OTUs were selected for downstream analysis, which accounts for more than 90 % of the whole dataset.

To identify the top 202 OTUs, DNA sequences of these OTUs, species of AOA and previously defined subclades of AOA (Jing et al., 2017) were codon aligned and used to construct a Maximum-Likelihood (ML) phylogenetic tree with MEGA 7.0 (Kumar et al., 2016). According to the result of model test (Bayesian Information Criterion calculation), the Tamura 3-parameter model, using discrete Gamma distribution with assumption that a certain portion of sites are evolutionarily invariables (T92+G+I), was used to construct the ML phylogenetic tree. The tree was further edited with iTOL (Letunic and Bork, 2016), with the relative abundances of top 202 OTUs displayed. The OTUs fell into the same subclades were further hierarchically clustered into ESTUs using SIMPROF test (method.distance=Euclidean) of the package “clustsig” in R (Whitaker and Christman, 2010), in which the OTUs having significantly similar distributional patterns (*P* value < 0.05) were grouped into same clusters (i.e. ESTUs).

### Statistical analyses

Non-metric multidimensional scaling (NMDS) plots were plotted using Primer 5 (Clarke and Gorley, 2001), to visualise the relationship between the AOA communities recovered in the 35 metagenomics datasets, based on the Bray-Curtis dissimilarity matrix. To characterize the ESTUs with environmental factors, correlation between the relative abundances of the ESTUs and environmental factors were analysed with Pearson test using package “Hmisc” in R; and the correlation plots were generated with the package “Corrplot” in R (Wei and Simko, 2013). The environmental factors were downloaded from the www.pangaea.de/.

## Results and Discussions

### High diversity of AOA in the metagenomics datasets

The top 202 OTUs were all affiliated to WCA, WCB and SCM-like clades, except two unclassified OTUs (OTU 53 and 154) (Fig. 2). Using the sequences of previously defined subclades of WCA and WCB (Jing et al., 2017) as references, all the subclades were detected in the *Tara* Oceans metagenomics datasets. In addition to that, new subclades of WCA (WCAIV) and WCB (WCAV) were identified (Fig. 2). The previously defined WCBII (Jing et al., 2017) was further divided into WCBII and WCBIII; and WCBIII in previous study (Jing et al., 2017) was renamed as WCBIV in this study (Fig. 2). Although there were 10 subclades of AOA commonly detected in the oceans (Fig. 2), only WCAI and SCM1-like has their representative cultures, *N. brevis* CN 25 (Santoro et al., 2015) and *N. maritimus* SCM-1 (Könneke et al., 2005) (Fig. 2). Based on the result of SIMPROF test, the OTUs of most subclades were grouped into ESTUs, except WCAIV, WCBII and WCBV, in which the OTUs showed homogenous distributional patterns (Fig. 2). Besides that, the genetically similar OTUs were not necessary to be included in the same ESTUs. For examples, within WCAI, the OTUs within WCAI-C were genetically similar, while the OTUs of WCAI-B were not closely clustering together in the phylogenetic tree (Fig. 2).

**Fig. 2.**
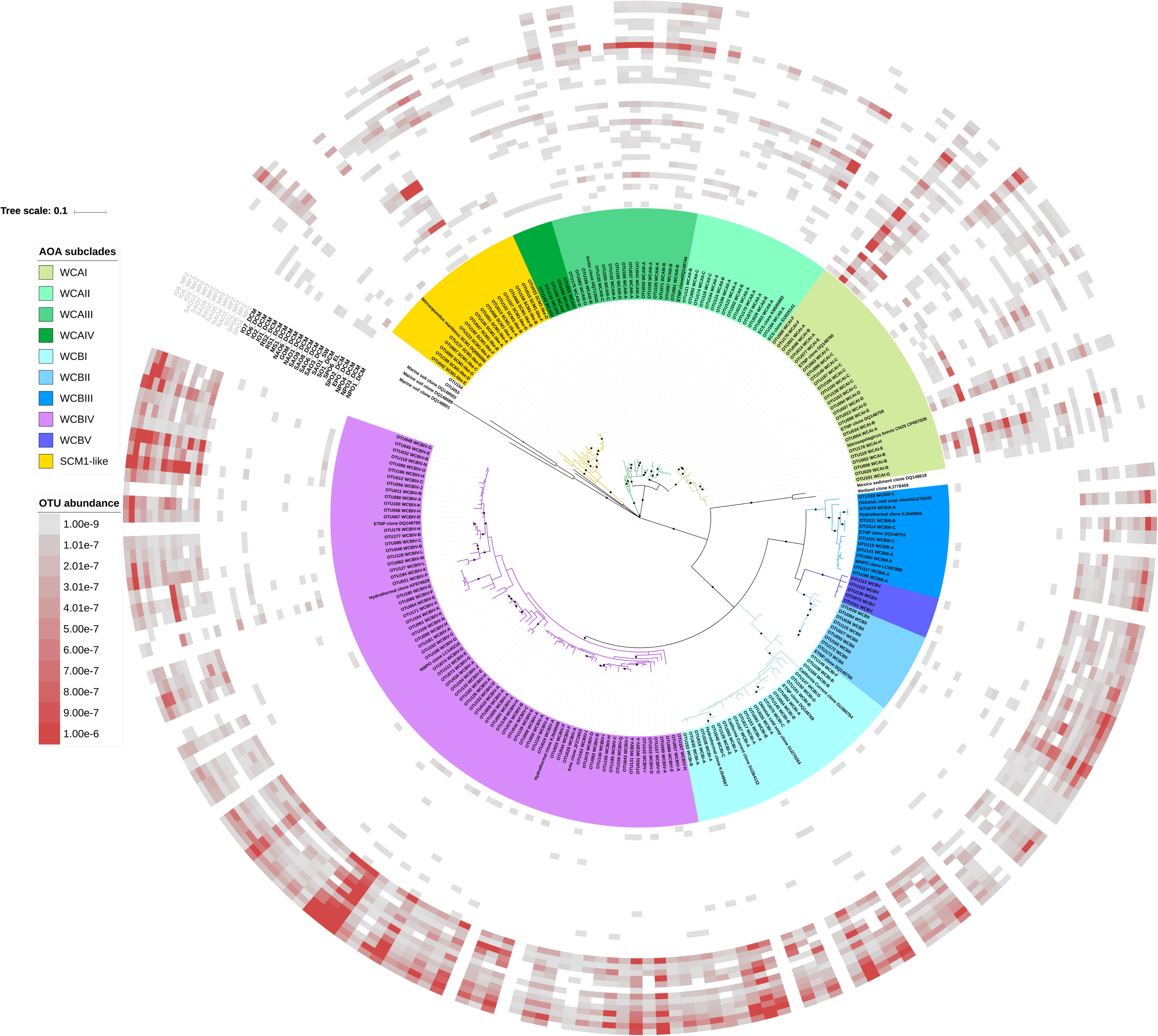
Maximum likelihood phylogenetic tree constructed using DNA sequences of top 202 OTUs, reference sequences of marine AOA cultures and previous defined subclades of WCA and WCB. The bootstraps test was conducted for 1000 times, and the values which higher than 60 % were displayed as black circles. The OTU abundances referred to that OTU affiliated reads divided by total clean reads in particular metagenomics datasets. The OTUs were labelled with their corresponding ESTUs (e.g. WCAI-A, WCAI-B, etc).

Considering the high genetic diversity of WCA and WCB and that the previous studies quantified the whole group of WCA and WCB (Mosier and Francis, 2011; Smith et al., 2014; Smith et al., 2016; Jing et al., 2017; Santoro et al., 2017), the sequences of qPCR primer sets of WCA and WCB (Mosier and Francis, 2011) were compared with representative sequences of the top OTUs of all the subclades (S2 Table). The result showed that both primer sets were well designed and can cover all the subclades, except WCAII, which has two bases mismatched with the WCA-amoA-P (Taqman probe). It has been reported that mismatch between primer set and template may influence the efficiency of amplification (Liu et al., 2006; Stadhouders et al., 2010). Therefore, the current primer set of WCA (Mosier and Francis, 2011) may underestimate the abundance of WCA, when WCAII predominates the AOA community. The actual influence of the mismatches to detection of WCAII is waiting for validation in future study.

### Global distributional pattern of AOA clades / subclades

The AOA communities in the euphotic zones were predominated by WCA and SCM1-like, while the communities in MPZs were mainly predominated by WCB (Fig. 3). The general vertical distributional pattern of WCA and WCB in this study agreed with the previous findings (Beman et al., 2008; Sintes et al., 2013; Sintes et al., 2016; Jing et al., 2017; Santoro et al., 2017). At the level of subclade, WCAI was the most dominant subclade in the euphotic layers of open oceans, followed by WCAII and WCAIII (Fig. 3). However, in the Gulf of Mexico and Red sea, WCAII became more dominant than WCAI. Previous study in the western North Pacific Ocean suggested that WCAII was specific to the Western Pacific Ocean (Jing et al., 2017), because the WCAII affiliated sequences in the NCBI nucleotide database were all recovered from the East China Sea (Hu et al., 2011). Our result suggested that WCAII is indeed globally distributed, however, it is predominant in the ecosystems of marginal seas. SCM1-like was another major member of the AOA community in the euphotic zone, which mainly predominated the waters above the latitude of 35° in both hemispheres. This agreed with previous study that *N. maritimus* SCM-1 affiliated sequences were detected in Arctic Ocean and Antarctic coastal waters (Kalanetra et al., 2009). Our results suggested niche separation of WCAI, WCAII and SCM1-like in the euphotic zones of the oceans, which are predominant in open oceans, marginal seas and high latitude waters, respectively.

**Fig. 3.**
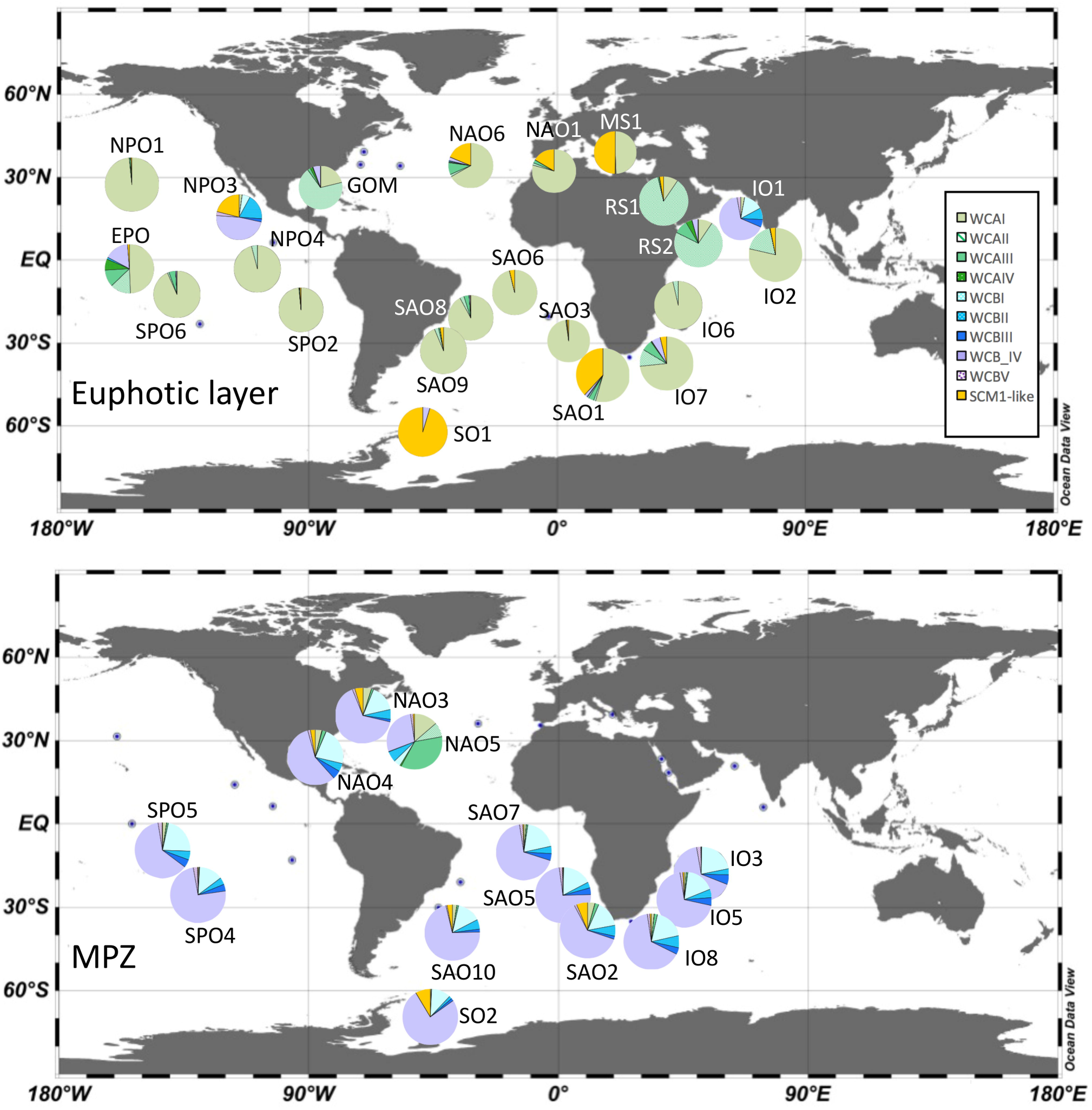
Distributional pattern of AOA communities in the global oceans. The community structures were consisted of the relative abundances of top 202 OTUs.

In the MPZs, WCBIV is the most dominant subclades, followed by WCBI, WCBII and WCBIII. However, it should be noticed that WCA and SCM1-like was also detected in the MPZs. Although the relative abundances of these two clades in the MPZ AOA communities were low (Fig. 3), their significant relative abundances in the MPZ metagenomics datasets suggested that they are more evenly and widely distributed than WCB (Fig. 2). The vertical profiles of WCA abundance have been analysed in several studies (Sintes et al., 2016; Smith et al., 2016; Jing et al., 2017; Santoro et al., 2017), which showed different results. In the equatorial Pacific and the northeast Pacific Ocean, the abundance of WCA decreased dramatically in the upper MPZ (near zero) (Santoro et al., 2017) and lower MPZ (10^2^ gene copy per liter) (Smith et al., 2016), respectively. However, at some stations in Atlantic Ocean and northwest Pacific Ocean, the abundances of WCA remained significant in the MPZs (10^3^ – 10^4^ gene copies per liter) and were comparable with that in the euphotic zones (Sintes et al., 2016; Jing et al., 2017). Together with our results (Fig. 2), it is not necessary for the distribution of WCA to be restricted in the euphotic zone and upper MPZs, and the significance of WCA and SCM1-like in the MPZs and even deeper water (Jing et al., 2017) is worthy for further exploration.

### Vertical segregation of AOA communities

At 7 geographic locations, AOA communities were recovered in both euphotic zones and MPZs (Fig. 4). The communities in the euphotic layers (IO6, IO7, SAO3, SAO6, SAO9, SPO6 and SO1) were more heterogeneous than those in the MPZs (IO5, IO8, SAO2, SAO7, SAO10, SPO5 and SO2) (Fig. 4). Especially in the Southern Ocean, the community in the euphotic zone (SO1) was distant from all other AOA communities in the dataset, while community in the MPZ (SO2) still clustered closely with other MPZ communities (Fig. 4). Similar patterns of bacteria and archaea communities have been reported in Pacific Ocean, in which the communities in surface waters are more heterogeneous than in the deep water (Jing et al., 2013; Xia et al., 2017). It was explained that the communities are less stable under the fluctuating physiochemical factors in the surface waters (Jing et al., 2013; Bryant et al., 2016). Except NAO5, which was sampled in upper MPZ (246 m in depth), all the MPZs datasets were originated from the samples collected in lower MPZs (> 500 m in depth) (S1 Table). Although the AOA communities were located in the lower MPZs in different oceans, they were all clustered together closely (Fig. 4). This is because the relative abundances of WCB subclades in the AOA communities were similar among the lower MPZs of different Oceans (Fig. 3), which can be explained by that the relatively stable environmental conditions in deep waters select the similar community of prokaryotes (Jing et al., 2013; Xia et al., 2017) and support the theory that “everything is in everywhere, but, the environment selects” (Baas-Becking, 1934).

**Fig. 4.**
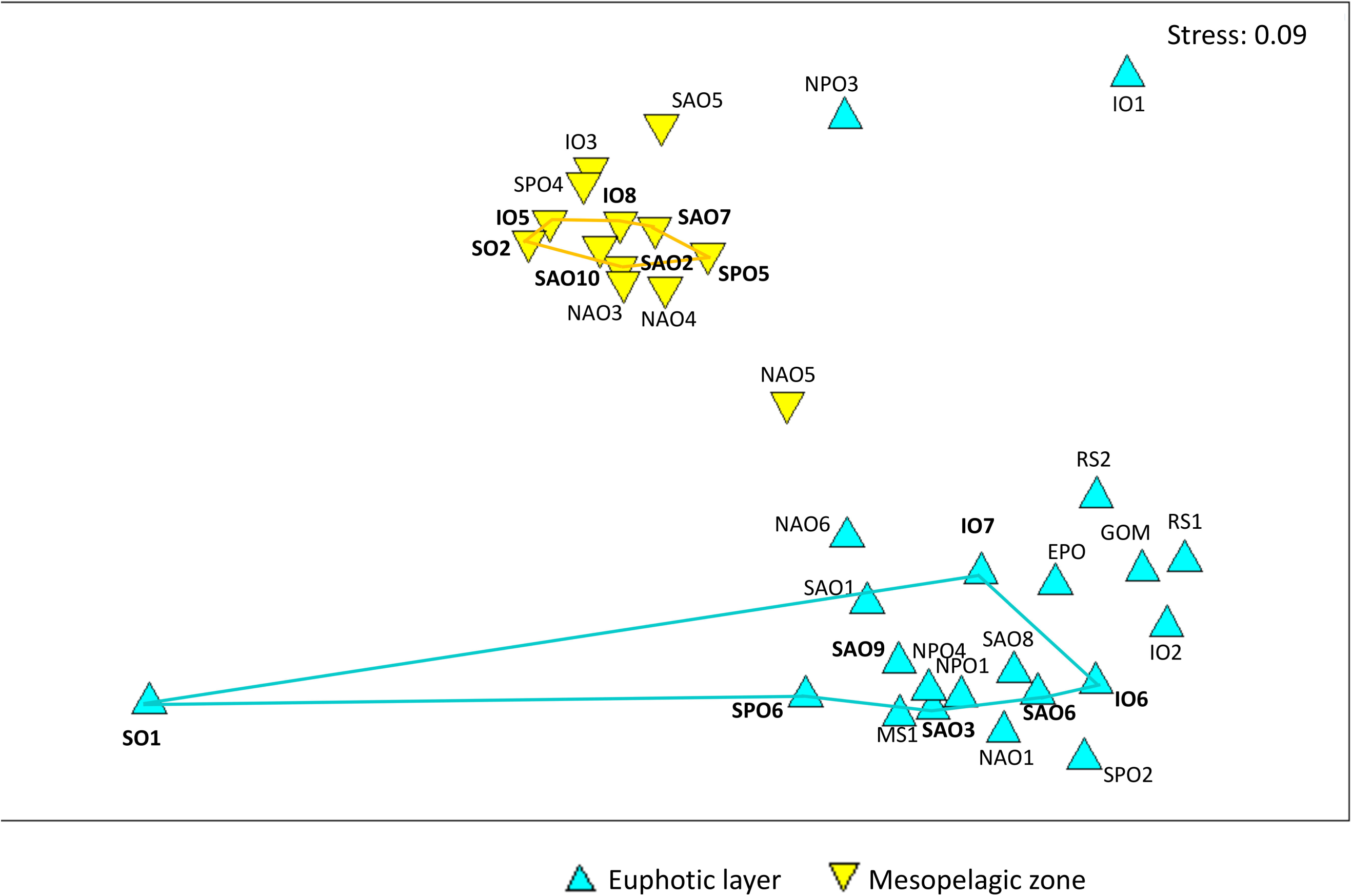
NMDS plot showed relationship between the AOA communities recovered from metagenomics datasets. The datasets that have corresponding datasets in euphotic zones or MPZs (same geographic locations) were bolded.

The vertical segregation of AOA communities was disrupted in the upwelling regions of northeast tropical Pacific Ocean (NPO3), Arabian Sea (IO1) and Equatorial Pacific Ocean (EPO) (Fig. 3, 4) (Wyrtki, 1967; Ulloa et al., 2012), where WCB were dominant in the euphotic layers (Fig. 3). Significant abundance of WCB was also detected in the shallow water of upwelling regions in Monterey Bay (Smith et al., 2014) and southeast Pacific (Molina et al., 2010). Our result showed that the vertical segregation of AOA communities is stable and predictable in the oceans, and physical force that transporting WCB to euphotic layers in upwelling region is the main factor of disrupting the segregation.

### Characterizing the ESTUs of AOA with environmental factors

For finer resolution of the connection between AOA genetic diversity and environmental factors, instead of subclades, the correlation between ESTUs of different subclades and environmental factors were examined using Pearson test. As the result, the ESTUs within same subclades could be correlated to different environmental factors (Fig. 5). Within WCAI subclade, three dominant ESTUs (WCAI-A, WCAI-B and WCAI-C) showed different correlations with depth. The WCAI-A prefers shallower water with low concentrations of organic nutrients WCAI-C prefers deeper waters with low temperature, while WCAI-B has no significant correlation with depth (Fig. 5). This agreed with the distribution patterns of these ESTUs that the relative abundance of OTU1 (WCAI-A) was higher in the euphotic zone, OTU46 of WCAI-B did not show obvious distributional pattern and OTU63 of WCAI-C was mainly distributed in the MPZs (Fig. 2). In previous studies which quantified the whole group of WCA, WCA was most abundant in the euphotic zone and remained significant abundance in the MPZs (Sintes et al., 2016; Jing et al., 2017). It can now be explained that the WCAI-A (top ESTUs of WCA) is responsible for the high abundance of WCA in the euphotic zone, while WCA in MPZ is mainly contributed by WCAI-C. The vertical succession between WCA and WCB along the water column has been well documented (Beman et al., 2008; Sintes et al., 2016; Smith et al., 2016; Jing et al., 2017; Santoro et al., 2017). In addition to that, our result suggested that vertical succession is also existing among the ESTUs of WCA. The ESTUs of WCAII have positive correlations with salinity and temperature (Fig. 5), because they were predominant in the low latitude marginal seas (Fig, 3), where the salinity and temperature were high (S1 Table). Similar to WCAI-C, WCAIII-A and WCAIII-C were also positively correlated with water depth. Besides the physical factors, a number of ESTUs of WCAI and WCAIII were positively correlated with nitrate : silicate ratio, which agreed with the recent finding that the abundance of WCA was positively correlated with this ratio in Equatorial Pacific upwelling region (Santoro et al., 2017). The nitrate : silicate ratio was treated as indicator of remineralization in their studied region (Raimbault et al., 1999; Jiang et al., 2003; Buesseler et al., 2008), and the positive correlation implied that WCA is a major player of nitrification and the trace metal and ammonium released during remineralization process are important requirement of WCA (Santoro et al., 2017). Our result showed significant and positive correlation of WCAI and WCAIII with nitrate : silicate ratio in the larger geographic scale. However, it should be noticed that the nitrate : silicate ratio may have other implications, which are depended on phytoplankton community structure and the geographic locations (Koike et al., 2001; Bibby and Moore, 2011). Therefore, further explorations are needed for verifying the relationship between WCA (or even WCA subclades), nitrate : silicate ratio and remineralization of organic matter.

**Fig. 5.**
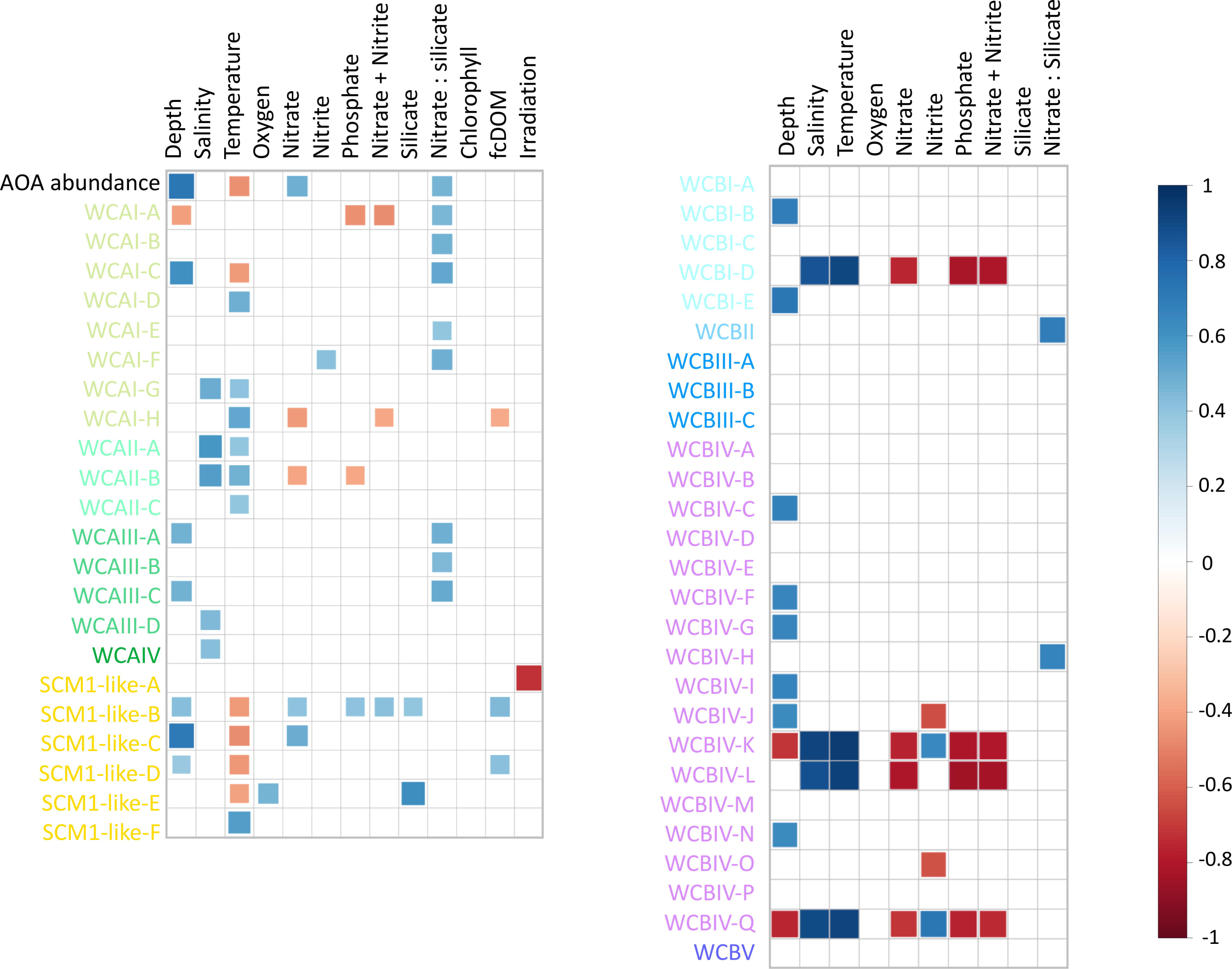
Correlation of relative abundances of total AOA and AOA ESTUs with environmental factors using Pearson test. The significant correlations (P-value < 0.05) were displayed. The ESTUs of different subclades were distinguished with different colors, which are identical to the coloring in Fig. 2 and 3.

The SCM1-like-A was negatively correlated with irradiation, while the rest ESTUs of SCM1-like clade did not (Fig. 5). SCM1-like-B, D and E were negatively correlated with temperature, agreeing with the general pattern that SCM1-like clade prefers high latitude waters (Fig. 3). Besides that, similar to WCAI-C and WCAIII, SCM1-like-C has a strong and positive correlation with depth (Fig. 5), which explains their main distribution in the MPZs (Fig. 2). Culture based study has discovered that different marine strains of *Nitrosopumilus* (SCM-1, PS0 and HCA1) had different sensitivity to photoinhibition (Qin et al., 2014), which supports our result about SCM1-like-A. Moreover, the positive correlations of SCM1-like-B and D with coloured dissolve organic matter (Fig. 5) also agreed with previous finding that some strains of *Nitrosopumilus* were obligate mixotrophs (Qin et al., 2014). The similar results of culture based study and our field study suggested that, the correlations between AOA ESTUs and environmental factors provides useful knowledge about the ecophysiology of AOA, especially the uncultured subclades.

WCB was rare and nearly undetected in the euphotic zone (Fig. 2), therefore, only the samples from lower MPZs (594 – 989 m in depth) were included in the correlation analysis. Some ESTUs of WCBI and IV showed preferences to deeper waters (Fig. 5). The WCBIV-K, L and Q showed similar pattern of correlation with salinity, temperature and macro-nutrients, which is because they were only found in rare number in the MPZ of the Southern Ocean (SO2), where was low in temperature and high in macronutrients (S1 table). Besides that, WCBII and WCIV-H showed positive correlations with nitrate : silicate ratio. Because the environmental condition in the MPZs is less variable, the ESTUs of WCB in the MPZs did not show significant correlations with most of the environmental factors (Fig. 5). Moreover, since the number of datasets from MPZs are limited, the correlation results of WCB may not be as significant as that of WCA and SCM1-like.

Based on our analysis, both subclades of the same clades and ESTUs of the same subclades could have different distributional patterns and correlation with different environmental factors (Fig. 2; 3 and 5), indicating the micro-diversity of AOA in the oceans should not be overlooked. In addition to studying the distribution of the whole group of WCA or WCB (Sintes et al., 2016; Smith et al., 2016; Jing et al., 2017; Santoro et al., 2017), analysing micro-diversity of WCA and WCB in the global scale refined our understanding to the ecophysiology of uncultivated AOA groups in marine ecosystems. Recent high-resolution genetic analysis of *Synechococcus* and unicellular cyanobacteria diazotroph (UCYN-A) also showed that the sublineages or ESTUs have different distributional patterns due to their different preference and sensitivity to certain environmental factors, causing niche separation in the oceans (Farrant et al., 2016; Cheung et al., 2017; Turk-Kubo et al., 2017).

Our study provided the first insight to the micro-diversity of AOA in the global ocean, and demonstrated the importance of high resolution genetic analysis of previous defined ecotypes. In addition to large geographic scale, high resolution studies in mesoscale regions with steep environmental gradients may provide more insights to the correlations between environmental factors and microorganisms (Robidart et al., 2014; Cheung et al., 2017). In addition to macro-nutrients, trace elements have also been suggested to influence the distribution of WCA and WCB (Santoro et al., 2017). For better understanding of the eco-physiology of uncultivated marine AOA, the abundance and high resolution distributional patterns of AOA subclades can be analysed using subclade specific qPCR primer sets in future studies.

## Acknowledgements

We thank the *Tara* Oceans for the huge open accessed metagenomics datasets of global oceans. This work was funded by the National Key Scientific Research Project (2015CB954003) sponsored by the Ministry of Science and Technology of the PRC and RGC GRF grant 16101917.

## Supplementary information

S1 Table. Information of the metagenomics datasets used in this study.

S2 Table. Mismatch between the qPCR primer sets that targeted WCA and WCB clades (Mosier and Francis, 2011) with the representative sequences of the AOA subclades.

